# Incorporating spatial diffusion into models of bursty stochastic transcription

**DOI:** 10.1101/2024.10.01.616185

**Authors:** Christopher E. Miles

**Affiliations:** Department of Mathematics, Center for Complex Biological Systems, University of California, Irvine

## Abstract

The dynamics of gene expression are both stochastic and spatial at the molecular scale. Mechanistic models of mRNA count distributions have revealed countless insights but largely neglect the frontier of subcellular spatial resolution. The spatial distribution of mRNA encodes their dynamics, including inherently spatial processes like transport to the nuclear boundary for export. Due to the technical challenges of spatial stochastic processes, tools for studying these subcellular spatial patterns are still limited. Here, we introduce a spatial stochastic model of nuclear mRNA with telegraph transcriptional dynamics. Observations of the model can be concisely described as following a spatial Cox process driven by a stochastically switching partial differential equation (PDE). We derive analytical solutions for spatial and demographic moments and validate them with simulations. We show that the distribution of mRNA counts can be accurately approximated by a Poisson-Beta distribution with tractable parameters, even with complex spatial dynamics. This observation allows for efficient parameter inference demonstrated on synthetic data. Altogether, our work adds progress toward a new frontier of subcellular spatial resolution in inferring the dynamics of gene expression from static snapshot data.

## I. INTRODUCTION

Gene expression at the molecular scale is stochastic [1, 2]. Consequences of this variability span development and disease [3–5]. Over the past decades, a vast body of research has evolved on constructing and analyzing increasingly intricate biophysical models to disentangle the sources and functions of gene expression stochasticity [6–13]. More recently, these mechanistic models have also seen uptake and success in revealing insights from mRNA count distributions from imaging and sequencing transcriptomics technologies [14–21].

Intertwined with stochasticity, gene expression is also an inherently spatial process at the subcellular scale [22]. After transcription at distinct locations within the highly structured nucleus, mRNA must be transported through the nuclear interior to be exported through nuclear pores [23]. This spatial transport and export of mRNA into the cytoplasm is followed by translation into proteins and serves as a fundamental regulation of gene expression [24–27]. Imaging technologies now give access to observing these subcellular spatial processes at unprecedented resolution. For instance, single-molecule fluorescence in situ hybridization (smFISH) [28, 29] provides spatial locations of individual mRNA molecules within nuclei and cytoplasm for multiple genes [30, 31]. State-of-the-art analysis of this subcellular spatial data is largely phenomenological [32–34] and challenging to associate with biophysical mechanisms. Models that do mechanistically account for nuclear export largely do so by treating the nucleus as a homogeneous compartment [35–40] and struggle to incorporate fine-grained spatial features that influence mRNA dynamics. For instance, nuclear geometries and transcription site locations vary per cell even for the same gene [41] and shape the timescale of mRNA export. Perhaps even more importantly, spatial locations of mRNA encode the underlying dynamics of their production and degradation, information that is discarded by considering only counts. In sum, important subcellular spatial details are readily available from imaging, but the ability to incorporate them into current mechanistic modeling machinery is lacking. Motivated by this gap between data and theory, this work pursues the advancement of stochastic models of gene expression to the next frontier of subcellular spatial resolution.

The slow progress toward faithful subcellular spatial models of gene expression is an outcome of the immense challenges involved. Beyond the staggering complexities of the spatial organization of the nucleus, even simple spatial stochastic models have considerable technical obstacles facing their uptake [42]. For nonspatial models, the pursuit is a scalar stochastic quantity described by a distribution. The addition of space increases the complexity dramatically with both stochastic numbers and spatial locations of interest. From a computational perspective, this hurdle cannot be overstated. For instance, consider a 2D smFISH image discretized into *N × N* pixels. There is a temptation to employ any of the zoo of techniques that have enjoyed success for nonspatial models, including generating functions [43], finite-state projections [16, 44], or neural networks [45, 46]. However, one would seemingly need to perform these sometimes already costly or complex calculations for all pixels in the image, each with a distribution for counts. Even for a modest *N* ∼ 100, solving for *N* ^2^ distributions for a single image becomes computationally prohibitive. Therefore, the analysis of spatial stochastic models for gene expression must be carefully considered to ensure that it remains computationally tractable enough to associate with data.

To address the computational challenges of spatial stochasticity, we employ a spatial point process formulation of the problem [47]. Specifically, we build upon our recent work [48] that investigates inference of the dynamics in a model of nuclear mRNA that undergoes spatial stochastic birth, death, and diffusion. The resulting description is a spatial Poisson process with an intensity that is the solution of a deterministic partial differential equation. This description allows for the rapid evaluation of the likelihood that encodes both the stochastic number and positions of the particles simultaneously, including the important complication of heterogeneous domains [15, 41]. With knowledge of the diffusion coefficient (measurable through live-cell tracking [49]), both the birth rate *λ* and death rate *γ* can be recovered from steady state snapshot spatial data. This is in contrast to the nonspatial birth-death process, where only the ratio *λ/γ* can be recovered, highlighting the value of spatial information even for purely demographic inquiries.

The mathematics of our previous work hinges on a Poissonian birth process corresponding to a constitutive gene. The constitutive model neglects the bursty behavior of transcription and consequently fails to reconcile the dispersion (variance relative to the mean) of RNA counts seen in real data [50–52]. In this work, we extend the point process framework to a more realistic telegraph model [53] of transcriptional activity that stochastically switches on and off. The introduction of particle correlations creates a technical obstacle unaddressable with the previous work’s machinery. Here, we follow [47] and leverage the spatial Poisson representation of the process [54, 55] that yields a stochastic PDE for the intensity of a Poisson process, making it a Cox process. Motivated by the success of the Poisson representation in deriving valuable analytical results for nonspatial models [56–59], the focus of this work is interrogating exact and approximate analytical results on this SPDE-driven point process model for spatial bursty gene expression.

The outline of the paper is as follows. We first introduce the formulation of the spatial stochastic model of nuclear mRNA with stochastically switching transcription. Using the spatial Poisson representation, we show that observations of the model follow a Cox process whose intensity is described by a stochastically switching PDE. For a 1D model, we analytically compute spatial and demographic moments for the point process, including verification by comparison with stochastic simulations. The value of these moments is limited, but we identify that the full distribution of counts can be well-approximated by a Poisson-Beta distribution with analytically tractable parameters that encode the spatial effects. This distributional approximation is shown to extend to more realistic variations of the model, including a semi-reflecting boundary that approximates pores on the nuclear surface, and more realistic cell shapes with heterogeneous interiors. The results culminate into a proof-of-concept demonstration of inference on synthetic data with heterogeneous cell shapes and gene locations. Altogether, the work paves a clearer path toward mechanistic model-based inference of stochastic gene expression that faithfully incorporates subcellular spatial dynamics.

## II. MODEL

The model considered here is a spatial stochastic description of individual molecules of mRNA in the nucleus of a cell, shown schematically in Fig. 1. The nuclear geometry is encoded in a domain Ω with boundary ∂Ω. The mRNA molecules are created via transcription at a fixed gene spatial location *z* at a stochastically varying rate based on a discrete promoter state. The transcription rate follows a dichotomous noise process [60], switching between an “on” rate *λ* and an “off” leaky rate *λ*_leak_, at rates *α, β* respectively. Once transcribed, mRNA diffuses in the nuclear geometry with diffusivity *D* until they face two possible outcomes: degradation at rate *γ* or exportation at the nuclear boundary. Nuclear export rates are encoded in a parameter *κ* discussed in further detail later. Heuristically, *κ* = 0 corresponds to no exportation (a purely reflecting boundary) and *κ* = ∞ describes the instantaneous export of RNA at the nuclear boundary (purely absorbing). The model observations correspond to the spatial positions of the RNA *x*_1_, …, *x*_*n*_ within the nucleus, where both positions and numbers are stochastically evolving. Throughout, only steady state dynamics *t →* ∞ are considered, acknowledging the neglect of transient effects from cell-cycle dynamics [13, 39, 61]. For illustrative purposes throughout the manuscript, the domain is considered to be one spatial dimension with Ω = [− *R, R*], with *R* crudely interpreted as the nuclear radius.

**FIG. 1.**
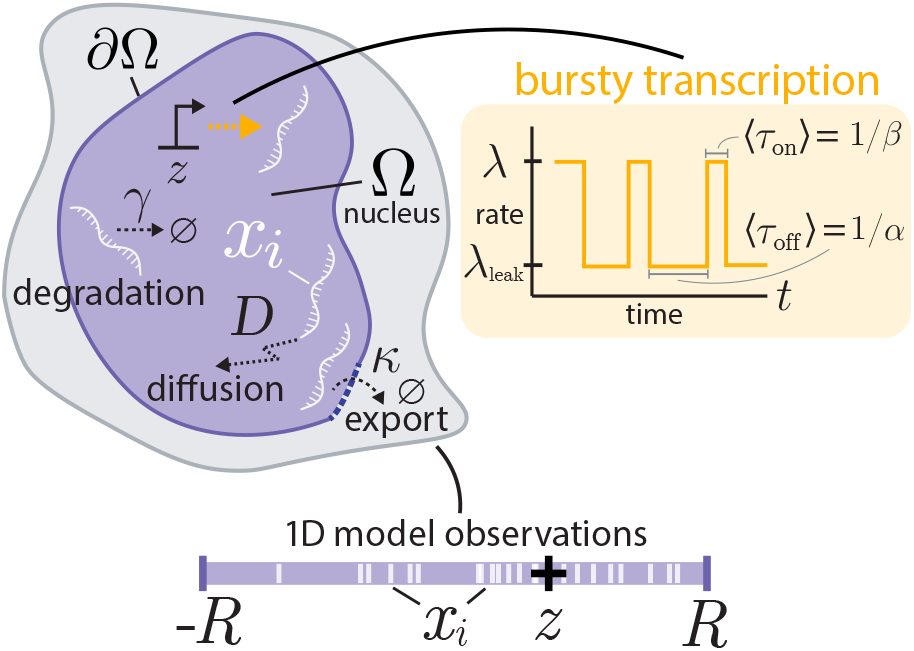
Schematic of the nuclear mRNA model. Transcription occurs at a spatial location *z* at a rate that stochastically switches between *λ* and *λ*_leak_ at rates *α, β*. After transcription, mRNA diffusive with diffusivity *D* until degradation at rate *γ* or export at the boundary, controlled by parameter *κ*. Throughout the manuscript, we primarily consider a 1-dimensional spatial model with Ω = [−*R, R*].

The rest of the manuscript pursues statistical descriptions of observations from this model. We reiterate the challenges for emphasis. Both the number and locations of the molecules are stochastic. Moreover, the temporally correlated nature of the birth process induces correlations between the molecules, so they may not be considered independently. These challenges are alleviated by the discovery that the model enjoys a straightforward spatial Poisson representation [47, 54, 55, 62]. With more details shown in Appendix A, we show that the particle locations, follow the spatial Poisson process

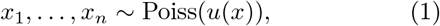

where *u*(*x*) corresponds to the steady state distributional solution *u*(*x, t*) of the stochastic PDE

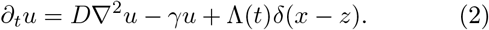

Λ(*t*) is the continuous-time (asymmetric) dichotomous noise process, switching between values {*λ*_leak_, *λ*}, at rates {*α, β*}, summarized by

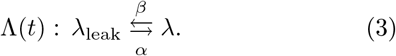

The stochastic PDE (2) can be written as a stochastically switching PDE

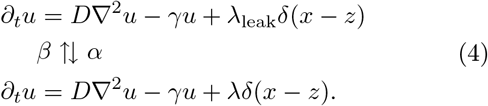

The stochastic nature of the intensity *u*(*x, t*) in the Poisson process (1) makes the observations *x*_1_, …, *x*_*n*_ a spatial Cox process, a doubly-stochastic Poisson process. We defer a discussion of boundary conditions for now.

It is worthwhile to note the contrast to the purely Poissonian birth (constitutive gene) case. If transcripts are produced at a constant rate *λ*, the Poisson process (1) remains, but now with a purely deterministic intensity

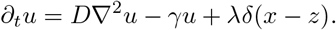

In this Poissonian birth scenario, the particles are entirely independent from each other. Therefore, the stochastic intensity underlying the point process can be attributed to arising from the correlated nature of the particles.

The Poisson representation provides clarity in encoding the locations and stochastic counts concisely. However, the resulting Poisson point process with an intensity driven by a stochastic PDE is not immediately illuminating. This leads us to pursue calculating emergent properties from this formulation.

## III. MODEL ANALYSIS

### A. Mean behavior

Our starting point for analysis of the model is in computing moments for both spatial positions and molecular counts. The mean behavior is straightforward to compute. Taking averages of (2) with respect to the realizations of Λ(*t*) gives the deterministic PDE for the average ⟨*u*(*x, t*)⟩

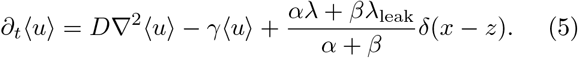

The mean of the counts for the Cox process (1) is the integrated mean intensity

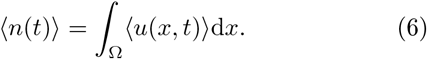

For ease of digesting the resulting formulae, we take *λ*_leak_ = 0, *κ* = ∞, and Ω = [− *R, R*] for now. In the steady state limit *t to*, (5) becomes the boundary value problem

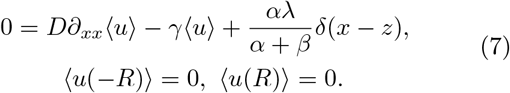

Calling *ρ* = *α/*(*α* + *β*), the fraction of time the transcription activity is on, the solution of (7) is

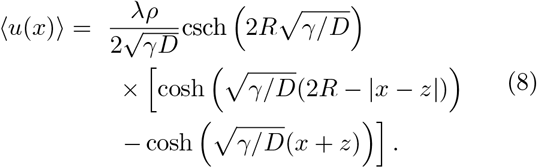

By (6), the total average number of molecules can then be computed by integrating (8)

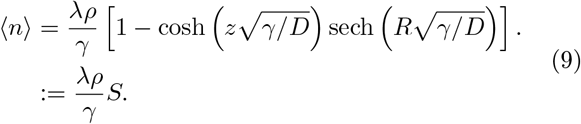

The resulting mean (9) deserves interpretation. The dimensionless scale factor *S* ∈ [0, 1] reflects the level of export out of the boundaries. As *S →* 1, the purely non-spatial mean is recovered and space plays no role in the molecular counts. Otherwise, *S* decreases with any factor that increases the overall flux out of the boundary: faster diffusion, smaller domains, or *z* closer to the boundary. *S* is a increasing function of *γ*, suggesting an interpretation of *S* as the probability a molecule is exported *before* it is degraded.

Verification of the predicted means (8) and (9) can be seen in Fig. 2. In panel a, the mean spatial position is shown for varying gene site location *z*, and all other parameters are fixed. As the gene site shifts closer to the boundary, the overall intensity level decreases. This is further highlighted in panel b, where the average total number of molecules goes to zero as the gene site approaches the boundary. The mean ⟨*n*⟩ also decreases with the degradation rate *γ* and diffusivity *D*, but increases with *α*. The predicted means and stochastic simulations show near-perfect agreement. Further details on the stochastic simulation can be found in Appendix B.

**FIG. 2.**
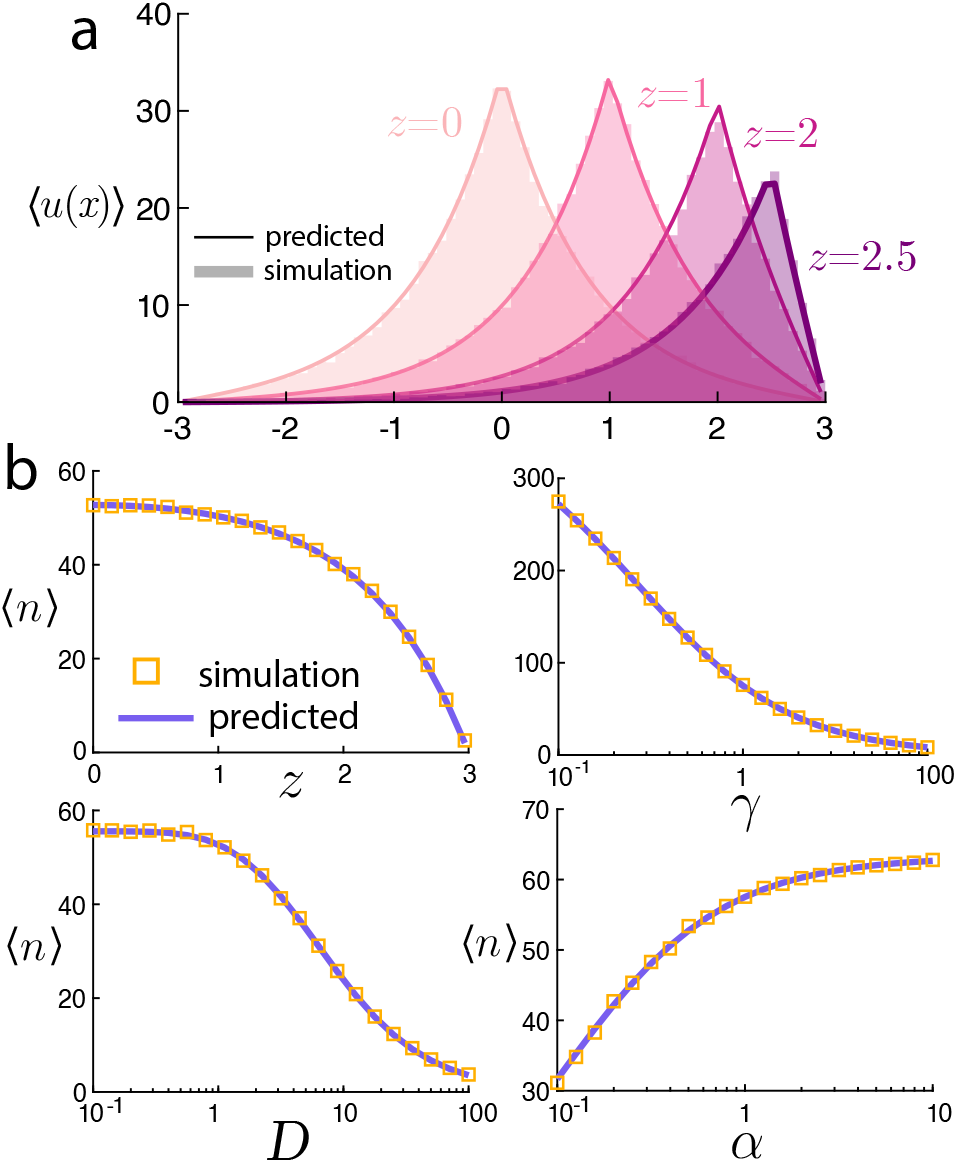
Mean steady-state behavior of the model. **a:** Mean positions ⟨*u*(*x*)⟩ from (8) varying source locations *z*. **b:** Mean number of particles ⟨*n*⟩ from (9) for various parameter sweeps. Simulations and predicted values closely agree.

### B. Variance of the molecular counts

The mean behavior of the telegraph model is effectively indistinguishable from Poissonian production with the lumped parameter *λp* as the effective production rate. We anticipate that higher-order moments do not bear this equivalence.

The variance for the total number of particles of the Cox process (1) can be computed by

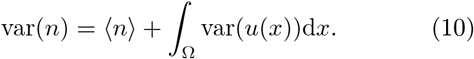

For the constitutive case, the intensity is deterministic, and var(*u*(*x*)) = 0 so that var(*n*) = ⟨*n*⟩ and a Poisson distribution is recovered. For the bursty process, *u*(*x, t*) evolves stochastically (2), so fluctuations in the intensity also manifest in contributing super-Poissonian variance to the molecular counts. The mean ⟨*n* ⟩ was computed in the previous section, so the determination of the variance of the counts is left to determine the variance of the intensity.

Contuining with purely absorbing boundaries *κ* =∞ in one spatial dimension Ω = [− *R, R*], we compute ∫ _Ω_ var(*u*)d*x* after a lengthy calculation. The result is a doubly infinite sum

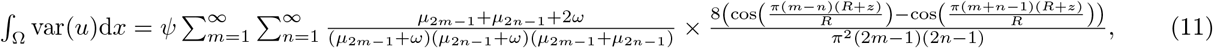

where *ψ* = *λ*^2^*αβ/*(*α* + *β*)^2^, *ω* = *α* + *β*, and *µ*_*m*_ := − *γ*− *π*^2^*Dm*^2^*/*(4*R*^2^). The calculation largely follows [63, 64] on the stochastic cable equation. The key observation is to note that the stochastic PDE solution to (2) can formally be written as

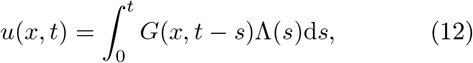

where G(*x, t*) is the appropriate Green’s function with a textbook series solution [65]. This Green’s function is combined with moments of dichotomous noise Λ(*t*) to yield this series solution for the integrated variance of the intensity. Further details of the calculation can be found in Appendix. C.

To the best of our knowledge and `Mathematica`’s abilities to simplify symbolic expressions, the infinite series (11) does not afford an elementary expression. It can be evaluated straightforwardly numerically. A comparison between the variance var(*n*) predicted by the series solution can be seen in Fig. 3. The parameters *D, γ, α* and *z* →*R* all yield a decrease in the variance of molecular counts. Furthermore, there is close agreement between the series prediction (11) and the stochastic simulations.

**FIG. 3.**
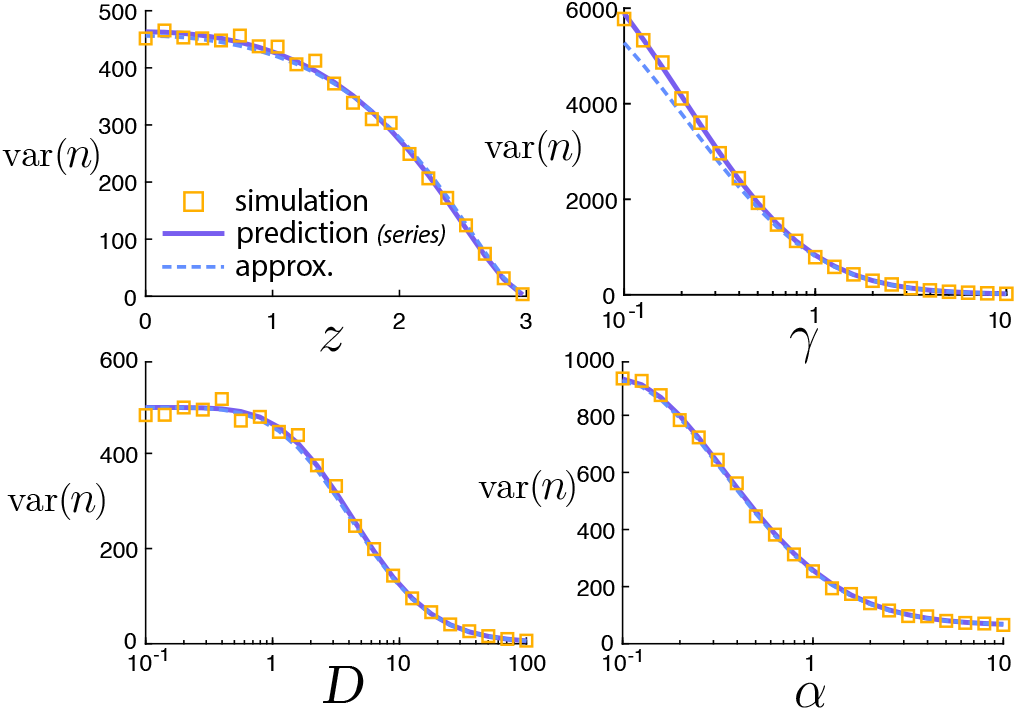
Variance behavior of the model. Variance of the number of molecules var(*n*) for various parameter sweeps. Simulations and predicted values from the infinite series solution (11) closely agree. Moreover, the approximate scaling value from the Poisson-beta distribution (14) is nearly (but not exactly) identical over the parameter ranges.

### C. Poisson-Beta distributional approximation

The difficulty in computing and digesting this series solution lends little hope to the direction of generalizing this machinery to more complex setups. Moreover, these moments are computed over realizations of the process in the same domain. Since the cell shape and gene site vary with each observation, it is not clear how moments may be directly connected to data with these heterogeneities. Instead, we identify a simple approximate description that will turn out to be surprisingly useful and accurate.

We motivate the approximation by reminding the reader that the mean molecular counts in (9) is *Sλα/*(*α*+ *β*), with the terms multiplying *S* interpreted as the non-spatial mean. One could arrive at this same answer by rescaling the parameters of the nonspatial process *λ*→*Sλ/γ, α*→*Sα/γ* and *β*→*Sβ/γ*. The division of *γ* arises from the lack of identifiability of a single parameter (time scale) in steady state. The factor of *S* will be the basis of the approximation.

The nonspatial variance is [7, 59]

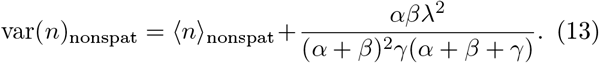

If we follow the same line of reasoning and rescale parameters *λ*→*Sλ/γ, α*→*Sα/γ* and *β*→*Sβ/γ* to account for spatial effects, this suggests

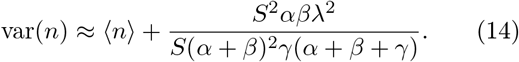

Values for this expression are plotted alongside the series solution in Fig. 3. Although this predicted value for the variance is distinct from the series prediction, the values are nearly indistinguishable over all parameters considered.

The nonspatial variance (13) corresponds to a Poisson-Beta distribution for the molecular counts, parameterized (15) by three values 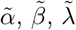

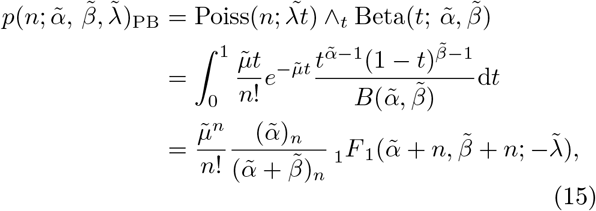

where ∧ _t_ denotes the mixture distribution with respect to *t*, _1_*F* _1_ is the confluent hypergeometric function, and (*c*)_*n*_ = *c*(*c* + 1) (*c* + *n −* 1) is the Pochhammer symbol.

The spatial variance approximation (14) is recovered with the choice of 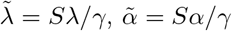, and 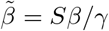Although this expression was obtained by heuristic scaling arguments of the moments, we can plot the full distribution with these parameter choices against the values from stochastic simulations. Fig. 4 shows these comparisons with remarkable agreement. In panel a we demonstrate the predicted distribution of *n* for the same setup as Fig. 2a and near-identical agreement is seen for all values of *n*.

**FIG. 4.**
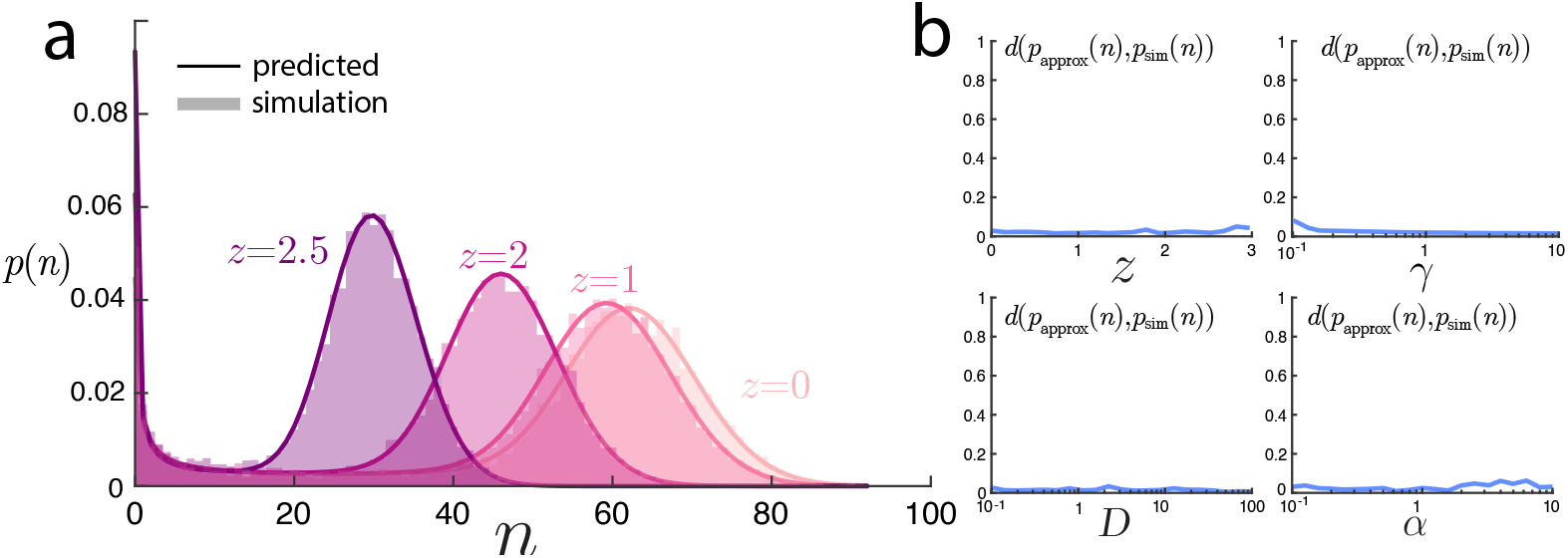
Comparison of the Poisson-Beta distributional predictions with simulations. **a:** For the same setups shown in 2a with varying *z*, the predicted full distribution of the number of molecules closely agrees with simulations. **b:** Over the parameter sweeps in Figs. 2 and 3, the Jensen-Shannon distance (between 0 and 1) between the predicted Poisson-Beta distribution of *n* and the values from simulations are consistently small, highlighting the broad applicability of the approximation.

One should be skeptical about the validity of this approximation. To investigate this, we computed the Jensen-Shannon divergence 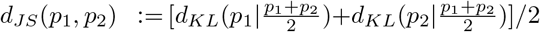 between the predicted Poisson-Beta distribution and the empirical distribution from simulations. This divergence takes values between 0 and 1, and across all parameter ranges tested shown in Fig. 4, the values were on the order of ≈ 0.01, suggesting remarkable agreement.

### D. Generalizing to semi-absorbing boundaries

The Poisson-Beta distribution provides an accurate prediction for the full distribution of molecular counts for purely absorbing boundaries and only requires a single deterministic PDE solution. To demonstrate this approach’s surprisingly broader applicability, we first extend the model to account for more realistic nuclear export. Consider a Robin boundary condition for the SPDE (2),

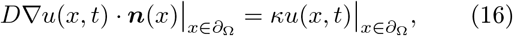

where ***n*** is the outward normal vector to the boundary ∂Ω. Such a boundary condition can arise from homogenizing the surface of the nucleus with absorbing pores [66, 67]. We interpret *κ* as controlling the kinetics of nuclear export [27]. With these boundary conditions, the PDE for the steady state mean intensity ⟨*u*(*x*) (7) ⟩ now becomes

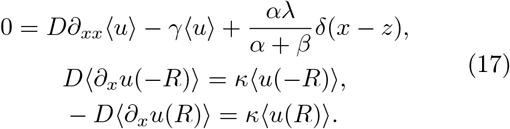

This can again be solved directly, and by (6) the mean number of molecules can be computed from the integral over the mean intensity, yielding a similar result to the *κ* = ∞ case (9)

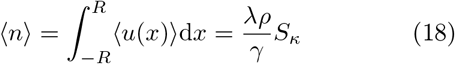

where *S*_*κ*_ is a slightly unwieldy but straightforward to compute expression

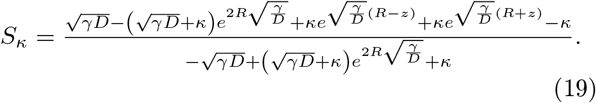

This scaling factor has the same interpretation: *S*_*κ*_ ∈ [0, 1] and can be understood as the probability of export before degradation. As *κ*→0, *S*_*κ*_→0, since no export occurs. In Fig. 5a and b, we show close agreement between stochastic simulations and the predicted mean behavior. As *κ* gets smaller, the number of molecules increases, and the intensity profile flattens out. In stochastic simulations, *κ* is interpreted in the partially-reflected sense [68], where larger *κ* encodes a higher probability of exit. Further simulation details can be found in Appendix B.

**FIG. 5.**
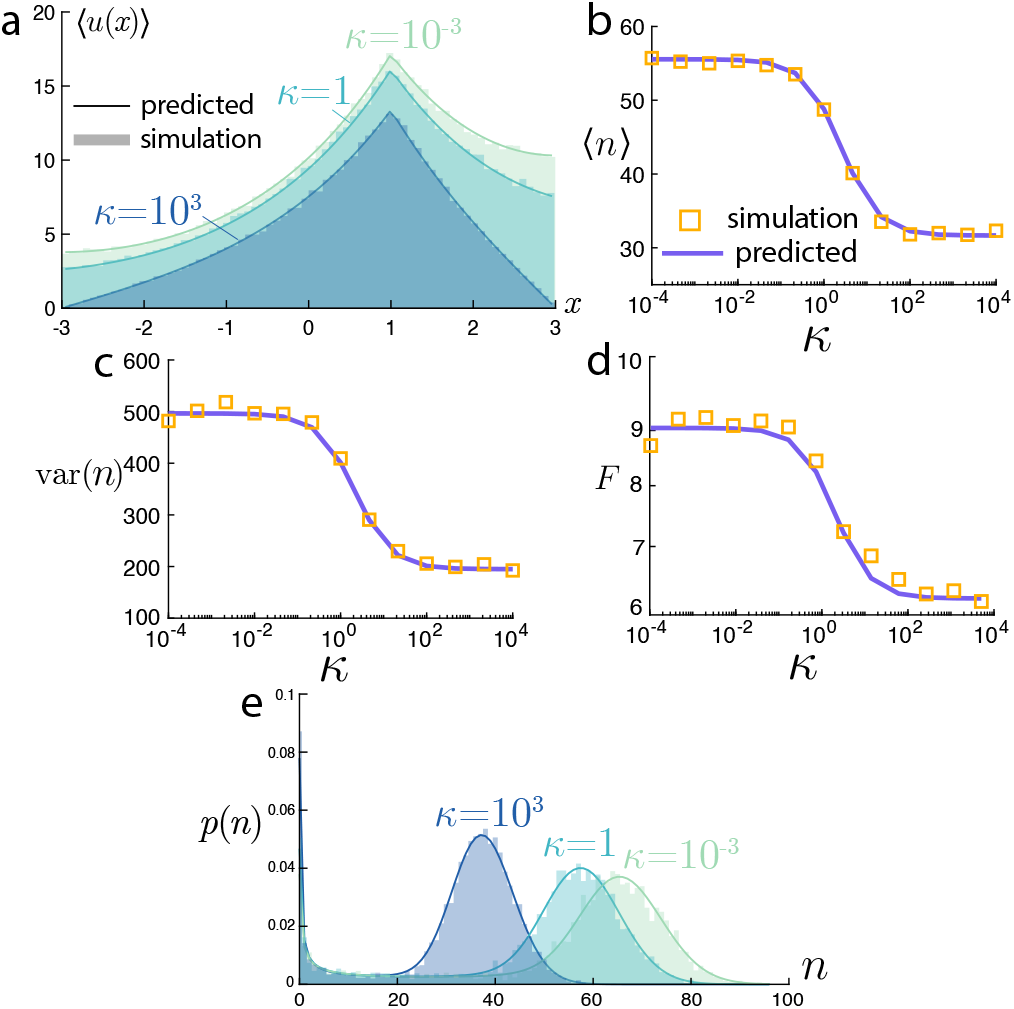
Influence of the export rate parameter *κ*. **a:** Mean positions ⟨*u*(*x*)⟩ for varying levels of *κ* from (17). **b:** Mean number from (18), **c:** variance, and **d:** Fano factor (variance/mean) of molecular counts *n* all decrease with *κ*. **e:** The Poisson-Beta predicted distribution shows close agreement with stochastic simulations for various *κ*.

We now consider the same parameter scaling for a Poisson-Beta distribution, 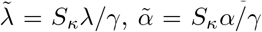, and 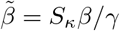. In Fig. 5c and d we show the predicted variance and Fano factor (variance divided by the mean). As *κ* increases, both the variance and Fano factor decrease, in agreement with experimental findings of [25] that show slowing down export (lower *κ*) leads to an increase in the nuclear mRNA Fano factor for several genes. Moreover, over this same range of *κ*, the Poisson-Beta prediction provides a remarkably accurate prediction for the full distribution of molecular counts, shown in Fig. 5e. This predicted distribution required only the computation of the deterministic PDE for the mean behavior (18). In contrast, the full variance calculation would have required tedious computations, ultimately likely to also result in an unwieldy infinite series solution. Thus, the finite *κ* scenario highlights the utility of the Poisson-Beta approximation.

### E. Fano factor interpretation of spatial effects

We are now equipped with the ability to study the nuclear mRNA model’s statistical behavior for various parameters and model variants. Before continuing to generalize the model, we take a brief interlude to highlight the value of an explicit spatial model in interpreting molecular counts of nuclear mRNA. In Fig. 6, we compare the Fano factor for the explicit spatial model

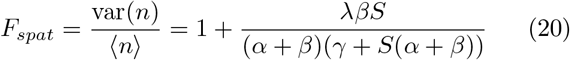

to the nonspatial variant neglects nuclear export

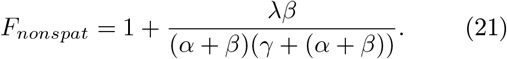

**FIG. 6.**
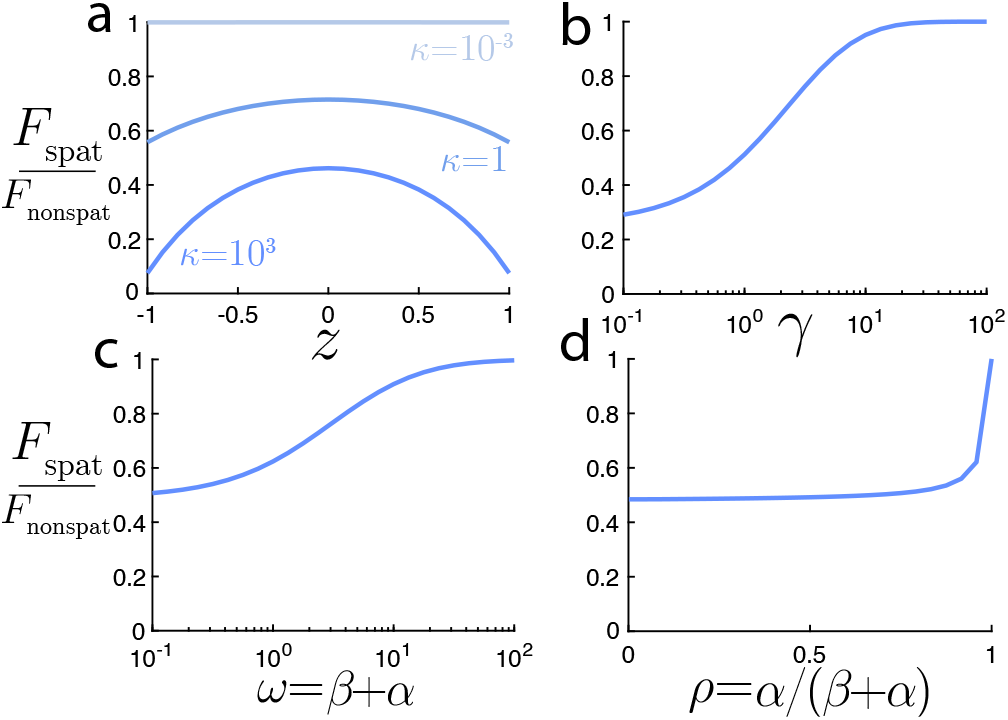
Comparison of the Fano factors for spatial and non-spatial models. **a:** Source locations with the largest *κ* and closest to the boundary have the largest deviation between spatial and nonspatial models. **b:** The degradation rate *γ* and **c:** transcriptional switching timescale *ω* both increase the agreement between the spatial and nonspatial Fano factors. **d:** The proportion of time in the “on” state, *p* has a relatively weak influence on the Fano factor ratio, except for nearly-constitutive *p* ≈ 1.

In Fig. 6a, we see that the largest deviation between the Fano factors of the spatial and nonspatial models occurs when exportation occurs most frequently: a gene site near the boundary (*z* ≈ R) and fast export (*κ* large). As the gene site moves to the interior of the nucleus or exportation slows, the two models more closely agree in their prediction. Notably, the Fano factor for the spatial model is always smaller than that of the nonspatial. Taken together, this can be straightforwardly understood by noting that export removes molecules from the count and therefore reduces the overall fluctuations. This point is supported in Fig. b, where larger values of the degradation rate *γ* are shown to lead to the smallest deviation between the spatial and nonspatial models. This is expected, as larger *γ* means the outcome of transcripts is dominated by degradation, and export plays less of a role in their dynamics. The interplay between export and the transcriptional state is less easy to predict. In Fig. 6c and d, we see that slow transcriptional dynamics *ω* = *α* + *β*, the overall switching rate leads to the biggest deviation. Moreover, the deviation is relatively robust to the fraction of time the transcription state is on, *p* = *α/*(*α* + *β*), except when *p ≈* 1 and production approaches a constitutive Poissonian process. Although these deviations between the nonspatial and spatial models may be easily understood and predicted, we emphasize their quantitative importance. If one were to fit a nonspatial telegraph model to the molecular counts with some Fano factor, the underlying mechanistic parameters would be incorrectly recovered and may lead to erroneous results.

### F. Two-dimensional cell with spatial heterogeneities

We make one last note about the applicability of findings toward more realistic setups. Undoubtedly, mRNA transport in the nucleus is not one dimensional, nor a spatially homogeneous process. The nucleus is crowded with various factors that are known to modulate mRNA motion [23, 49]. Moreover, the geometry of the nucleus itself plays a role in the export process because the mRNA must be transported to the boundary to be exported. In this last demonstration, we highlight the ability to handle these challenges within the currently presented framework.

In Fig. 7, we show a synthetically generated, two dimensional cell geometry. The nuclear interior is assumed to be heterogeneous, modeled by a spatially-dependent diffusion coefficient *D*(*x*). There are inherent technical challenges in interpreting spatially-dependent diffusivities [69, 70]. Here, we take an Itô interpretation with no justification beyond being the most straightforward to implement. In this vein, we also consider purely asboring boundaries (*κ* =∞). How can the distribution of mRNA counts and positions in this spatially heterogeneous domain be predicted? Based on the findings thus far, both of these distributions can seemingly be accurately approximately by a single deterministic PDE solve for the mean intensity. Motivated by this observation, we use `MATLAB`’s PDE Toolbox to solve the deterministic equation

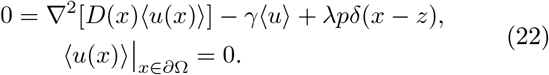

**FIG. 7.**
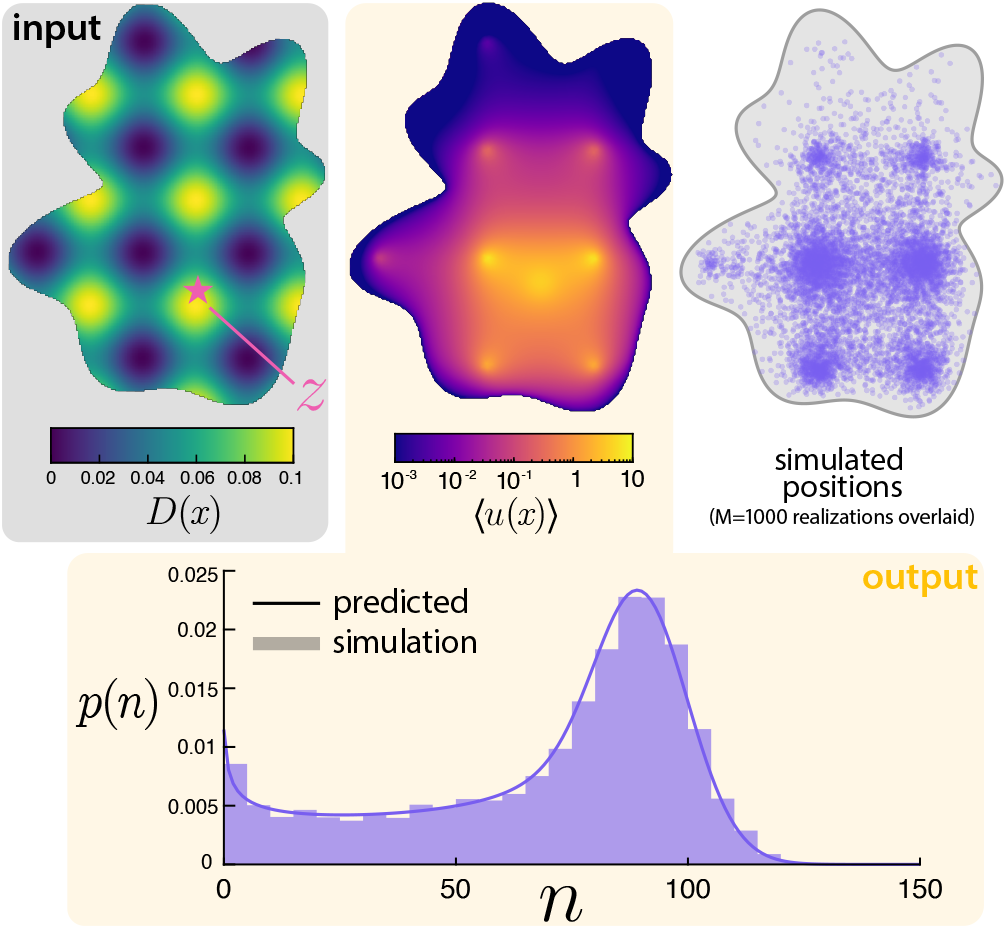
Generalizing the model to more realistic scenarios. The diffusivity *D*(*x*) is taken to be spatially varying (in the Itô sense) in a randomly generated two-dimensional domain. The mean number of particle ⟨*u*(*x*)⟩ is solved numerically from the PDE (22) and shown. The total mean is then used to specify the parameters of a Poisson-Beta distribution for the total number of particles *n*, which shows close agreement with stochastic simulations.

The choice of Itô interpretation manifests in the location of the derivative with respect to the diffusivity. The solution of this PDE for ⟨*u*(*x*) ⟩ is shown in Fig. 7 alongside spatial positions of several stochastic superimposed over several realizations. The prediction for positions and stochastic simulations demonstrate excellent agreement. Qualitatively, the molecules tend to get localized in regions of low diffusivity. Next, we turn to quantitative predictions. Motivated by the one-dimensional answers (9) and (18), we can define the analogous scale factor as

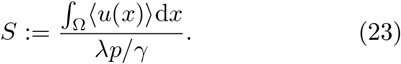

This integral can be computed from the numerical PDE solution. Then, we posit that *n* approximately follows a Poisson-Beta distribution with parameters 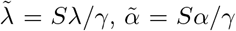, and 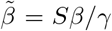. The resulting predicted distribution is shown in comparison against the counts from simulations in Fig. 7. This prediction using the numerical PDE solution for the scaling factor shows remarkable agreement with simulations. Although numerically solving a PDE on cellular geometries may be computationally costly, we emphasize that it seems far less costly than any current alternative approach for characterizing the distributions of both counts and positions. For instance, a generating function approach would seemingly require a PDE solution for each discretized pixel.

## IV. INFERENCE ON RATES WITH HETEROGENEOUS CELLS

In this last section, we outline a possible avenue to employing the findings in this work toward inference of model parameters with data. Although we have thus far achieved an understanding of forward predictions of the model, the Cox process observations provide distinct technical challenges in their inference. The likelihood for a single observation (cell) of the Cox process (1) with parameters ***θ*** is [71]

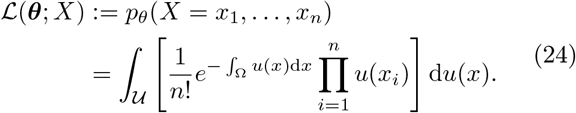

In other words, evaluation of the likelihood requires (infinite dimensional) integration over all possible realizations of the solution to the stochastic PDE (2). While there is extensive literature on sophisticated numerical approaches for performing inference with this Cox process likelihood [71], we instead focus on the possibility of a simpler approximation that leverages our earlier discussed findings.

As a motivating detour, we momentarily consider the likelihood of a simple Poisson process with deterministic intensity *u*(*x*) and underlying parameters ***θ***. In this case, the likelihood becomes the term inside the integral of the Cox process likelihood

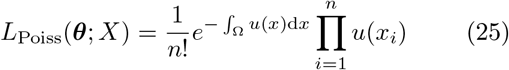

The observation we hope to emphasize can be seen by rearranging this likelihood and denoting ⟨*n*⟩ = ∫_Ω_ *u*(*x*)d*x* and *p*(*x*_*i*_) = *u*(*x*_*i*_)*/* ⟨*n*⟩. Then, the likelihood (25) can be written as

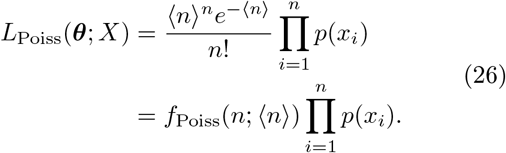

In other words, for the Poisson spatial point process, the likelihood can be decomposed into the contributions of the counts and positions. Motivated by this observation and the findings presented thus far, we approximate the full Cox process likelihood by

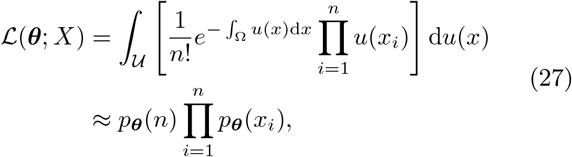

where the spatial distribution is computed from the expected positions

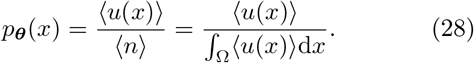

Aside from the motivating example with deterministic intensity *u*(*x*), this approximation does not seem to be exact. Instead, (27) can be interpreted as a “mean-field” approximation that does not fully account for correlations between counts and the spatial positions. Importantly, however, the quantities in this approximation are all tractable using the previously discussed results of this work. The mean behaviors used in (28) can be computed exactly. The count distributions *p*_***θ***_(*n*) were found to be well-approximated by a Poisson-Beta distribution with parameters computed from ⟨*n*⟩. Therefore, the evaluation of this approximate likelihood reduces to straight-forward analytical or numerical solutions to a single PDE for the mean intensity. In most scenarios we can imagine, this is far less costly than any Monte Carlo sampling technique for evaluating the true Cox process likelihood.

The obvious question that remains is whether this approximation to the likelihood is sufficiently accurate to do reliable inference. We address this question with a demonstration of inference on synthetically generated data, presented in Fig. 8. To mimic the realistic challenge of cell-to-cell heterogeneity, we generate *M* = 500 synthetic observations from the one-dimensional model, where each cell *i* has a randomly generated *z*_*i*_ and *R*_*i*_, but all kinetic parameters are fixed across the cells. We again assume steady-state conditions, so not all kinetic parameters are identifiable. We assume that the diffusion coefficient *D* may be measured through other means (for instance, live-cell tracking [49]), but the remaining parameters are unknown ***θ*** = [*λ, γ, α, β, κ*]. With this synthetic dataset, we employ the Cox process likelihood approximation (27) for maximum likelihood inference using `MATLAB`’s `fminsearch`. To simultaneously diagnose the identifiability of these parameters, we also compute the profile likelihood for each parameter, defined by

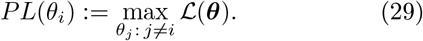

**FIG. 8.**
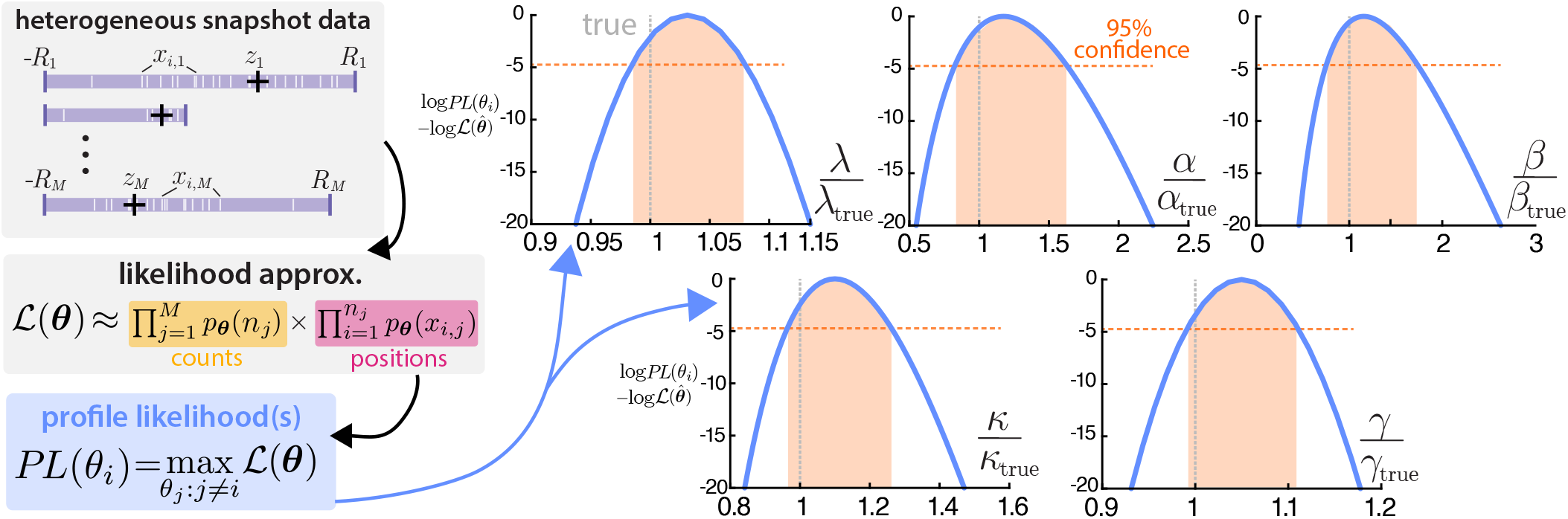
Demonstration of inference and identifiability on a heterogeneous dataset. A synthetic dataset of *M* = 500 images, each with randomly chosen *R* and *z* was generated. This is then used in an approximation of the full likelihood (27) from which profile likelihoods are computed. The profile likelihoods are plotted as *χ*(*θ*_i_) := log *PL*(*θ*_i_) − log ℒ (***θ***) to facilitate comparison with the threshold for an asymptotic confidence region for simultaneous inference of the parameters. Since all profile likelihoods decrease sufficiently fast over the windows of interest, structural and practical identifiable can be concluded.

That is, the parameter of interest is fixed to a specified value (determined by the functional input to the profile likelihood) and the remaining parameters are optimized. The profile likelihood is a standard diagnostic for identifiability of parameters [72]. The output of each profile can be compared to the maximizing value of the likelihood over some window of interest, often corresponding to a confidence region. If the profile likelihood takes on sufficiently distinct values over this window, the parameter can be interpreted as both structurally and practically identifiable [73].

In Fig. 8, we see that the maximizing values of the profile likelihoods closely agree with the true simulated parameters. Moreover, we consider a window corresponding to the asymptotic 95% confidence region for the recovery of all parameters with a threshold of *χ*^2^(0.95, 4), accounting for the 4 degrees of freedom in the optimization of each profile likelihood [73]. Over this confidence region, all 5 parameters appear to be structurally and practically identifiable. We remind the reader of the nonspatial variant, a Poisson-Beta distribution with only 3 identifiable parameters. It seems worth noting that all true parameters fall at the lower end of the confidence region, which may be indicative of biased estimates arising from this approximate inference procedure. We believe this proof-of-concept demonstration highlights how the findings of this work can be leveraged for inference, but optimization and dissection of this inference are left for the future.

## V. DISCUSSION

We have formulated and investigated a model of nuclear mRNA that explicitly incorporates nuclear spatial diffusion and telegraph transcriptional dynamics. The most fundamental finding is that observations of the model form a Cox process, a spatial point process with intensity corresponding to the solution of a stochastically switching PDE. This spatial telegraph PDE lends itself to some analytical tractability in one spatial dimension. The mean and variance of both spatial distributions and counts were computed and verified against stochastic simulation. The model predictions about the role of spatial processes qualitatively agree with experimental findings, most notably that slowing down nuclear export increases the dispersion of nuclear mRNA counts [25]. However, these basic calculations were unwieldy even for the simplest of the models, making quantitative comparison challenging. The main upside of our work is the observation that a Poisson-Beta distribution well-approximates the full count distribution and parameters of this distribution can be straightforwardly computed. This distributional approximation was shown to generalize broadly, including Robin boundaries that model nuclear export and generically shaped two-dimensional cells with spatially heterogeneous diffusion. The computational tractability of the count distribution empowers the ability to perform approximate inference. We show that with heterogeneous snapshots of cells with distinct sizes and gene sites, kinetic parameters are identifiable from the spatial distributions and counts. Altogether, our work paves fundamental theoretical progress toward connecting imaging data (for instance, from smFISH) of spatial distributions of nuclear mRNA to infer the spatiotemporal gene expression dynamics underlying them.

Our findings should not be viewed as in tension with work that models gene expression dynamics nonspatially, for instance, those that treat mRNA export as a unimolecular reaction between homogeneous nuclear and cytoplasmic compartments [36, 40]. Instead, our work sheds light on why these nonspatial models have found such success in their ability to fit observed RNA counts. Even with explicit spatial diffusion, nuclear export, and spatial heterogeneities, we found that a Poisson-Beta distribution, the same prediction as the nonspatial process, well-describes the count distribution across parameters. The observation that explicit diffusion until export in the nucleus can be modeled as a single-step reaction has been noted before [25], but we emphasize that care must be taken in interpreting the parameters. For instance, a decay term must be interpreted as a mix of degradation and spatial export, and our findings show how these terms are differently affected by spatial geometric factors. It seems challenging, perhaps impossible, to disentangle these effects through fitting to nonspatial models. In contrast, we have shown that fitting the distributional counts with spatial information empowers a new resolution of detail to interrogate the dynamics.

Commenting on the biological relevance of the work, we have not paid much attention to the choice of parameters used throughout the manuscript except in their use to highlight qualitative features. Although the nucleus is crowded and heterogeneous, approximately diffusive motion has been observed for some mRNA [49] with a diffusion coefficient on the order of *D* 0. ≈ 03um^2^/s [23]. Taken with a typical nuclear radius on the order of *R* ≈ 10um [29], this gives a diffusive timescale of *R*^2^*/D* ≈ 1hr. This is in agreement with the observation that export and degradation are on approximately the same time scales [25]. However, nuclear export is not instantaneous, nor occurs at every location on the nuclear surface [24, 27]. This suggests finite *κ* seems most appropriate, but we were unable to identify an approximate numerical value. It is unlikely that the full spatial distribution of diffusion coefficients can be identified as in Fig. 7. However, we believe this demonstrates the possibility of imaging nuclear condensates (nucleoli, speckles) that are known to mediate spatial organization of Mrna [23] and leveraging this spatial information in the modeling and inference process. There is some evidence that mRNA may be shielded from degradation prior to export [74], but degradation was included in the model for broad generality.

The telegraph process for transcriptional dynamics was chosen in part due to its popularity, but also as a minimally complex example that produces super-Poissonian dispersion of molecular counts. This dispersion is intimately linked with the correlation between the particles induced by the production process and causes the particle-wise machinery of our previous work [48] to fail. We anticipate this approach lends itself better to generalizing into other complexities of interest. In this vein, there are several avenues of promising future direction. We have considered only a single mRNA population, but one could imagine extending the framework to account for multiple species available from imaging such as multiple genes [31, 75] or both nuclear and cytoplasmic RNA [31, 40]. The telegraph transcriptional process also lends itself to generalization to more realistic multi-state production processes [76– feedback (with delay) [11, 81], or time-dependent cell-cycle dynamics [13, 61, 82]. Even more broadly, we hope the machinery of this work can be used to study other spatial regulations of gene expression [83, 84] that occur with distinct subcellular localization such as RNA splicing [29, 85] or phosphorylation [86], both of which

On the theory and computational side, the finding of a Poisson-Beta distribution for the counts was heuristically motivated. Due to the lack of proof, it remains unclear whether the distribution of counts genuinely follows this distribution or is merely well-approximated by it. We believed that this approximation corresponds to a truncation of the Green’s function series. However, the leading term of the series solution (11) does not seem to follow the form (14) conjectured from scaling arguments. Nonetheless, we believe this line of investigation is worthwhile to share and warrants further investigation in the future due to its computational tractability. Alternative computational approaches that seem likely to be fruitful include simulation-based inference [87, 88] and graph neural networks for stochastic reaction-diffusion that can generalize to arbitrary geometries [89].

## ACKNOWLEDGMENTS

The author was partially supported by NSF DMS CAREER-2339241. He is grateful for helpful discussions with Fangyuan Ding and Scott McKinley about this work.

## Appendix A: Poisson representation of the model

In this section, we provide further details on the claim in the main text that the observations of the discrete particle process follow a Cox process (1) with intensity governed by the stochastic PDE (2). The idea of spatial Poisson representations dates back to Gardiner [54, 55]. More recent work that is closely related and more accessible than Gardiner’s original exposition can be found in [47] and the geophysics literature [62]. We believe the explicit computation of this model with dichotomous noise has not been considered.

For simplicity that can be easily generalized, consider a 1D domain discretized into windows of size Δx, each with counts *n*_1_, …, *n*_*M*_. Denote the promoter state corresponding to Λ(*t*) in the main text as *j*(*t*) = {0, 1}. In state *j* = 0, transcription occurs at rate 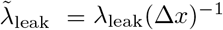 and in state *j* = 1, at rate 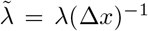. The gene site is fixed to be in the spatial location indexed *k*. In the discretized model, diffusion corresponds to a unimolecular reaction at rate 𝒟 = *D/*(Δx^2^).

Denote *p*_0_(***n***, *t*) = *p*(***n***, 0, *t*) and *p*_1_(***n***, *t*) = *p*(***n***, 1, *t*) respectively. The stochastic vector of counts ***n*** and promoter state *j* evolves via the Chapman-Kolmogorov equations

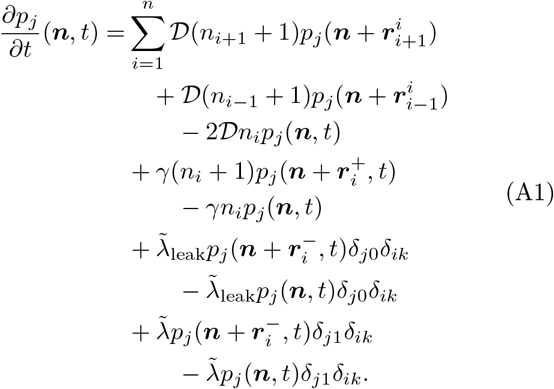

We have abbreviated the perturbation vectors 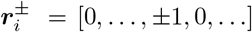 and 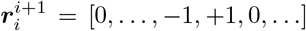. The Poisson representation takes the ansatz

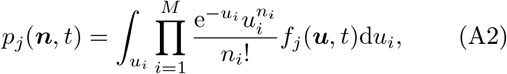

which gives the evolution of the intensities as

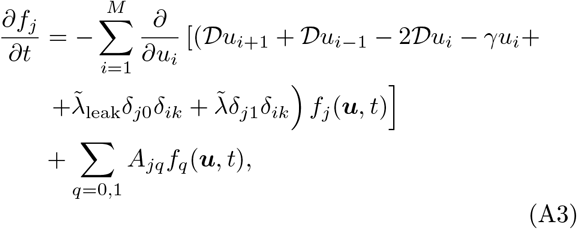

where *A* is the 2 × 2 generator matrix for the promoter switching process. This is the Chapman Kolmogorov equation for the piecewise deterministic Markov process (PDMP) system

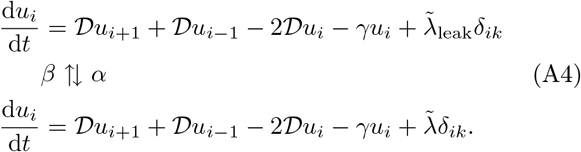

In the limit as Δx→0 with *u*(*x, t*) = *u*_*i*_(*t*)*/*Δx, the switching PDE (2) is recovered. We note the structural contrast of this result compared with related works [47, 62] that arrive at Gaussian SPDEs from biomolecular reactions or white-noise limits.

## Appendix B: Stochastic simulation details

For all stochastic simulations, the promoter state switching, birth of molecules, and their decay are handled by a standard Gillespie simulation [90] until some time *T*. To ensure steady state, *T* is chosen to be *T* = 100*/* min {*α, β, γ, λ, R*^2^*/D*}. For all particles living at the end of the simulation, their birth times are recorded. These candidate molecules are then simulated spatially, starting at their birth time until either *T* or they exit the domain. The final positions of all remaining molecules that have survived the Gillespie and spatial simulation steps are outputs of the simulation. In simulations with finite *κ*, the boundary is assumed to be a partially reflected diffusion [68]. If a particle Euler steps out of the domain, with probability *P* Δt it is absorbed. Based on that work, 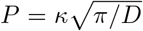 If the particle is not absorbed, it is then reflected. All finite *κ* simulations throughout the manuscript are in one spatial dimension, where reflection is straightforward. In simulations with heterogeneous *D*(*x*), the Euler step is interpreted in an Itô sense, 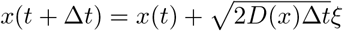.

### 1. Parameter values used in figures

Fig. 2: *z* = 0, *R* = 3, *α* = 0.5, *β* = 0.1, *λ* = 100, *λ*_leak_ = 0, *γ* = 1.5, *D* = 1. *N*_sims_ = 5000.

Fig. 3: *z* = 0, *R* = 3, *α* = 0.5, *β* = 0.1, *λ* = 100, *λ*_leak_ = 0, *γ* = 1.5, *D* = 1. *N*_sims_ = 5000.

Fig. 4: *z* = 0, *R* = 3, *α* = 0.5, *β* = 0.1, *λ* = 100, *λ*_leak_ = 0, *γ* = 1.5, *D* = 1. *N*_sims_ = 5000.

Fig. 5: *z* = 0, *R* = 3, *α* = 0.5, *β* = 0.1, *λ* = 100, *λ*_leak_ = 0, *γ* = 1.5, *D* = 5. *N*_sims_ = 5000.

Fig. 6: *z* = 0, *R* = 1, *α* = 0.5, *β* = 0.1, *λ* = 100, *λ*_leak_ = 0, *γ* = 0.75, *D* = 1, panels b-d: *κ* = 10^3^.

Fig. 7: *z* = [0, −0.5], ⟨*R*⟩ = 1, *α* = 0.75, *β* = 0.25, *λ* = 100, *λ*_leak_ = 0, *γ* = 1, *D*(*x, y*) = *D*_−_ + (D_+_ − *D*_−_)(cos(3π*x*) + cos(3*πy*) + 2)*/*4,, *D*_+_ = 0.1, *D*_−_ = 10^−3^, *N*_sims_ = 5000.

Fig. 8: Domains are randomly generated by *R*_*i*_ ∼ Γ(4, 1*/*4), *z*_*i*_ ∼ *β*(4*/*3, 4*/*3) × 2*R − R*, so ⟨*R*⟩ = 1, var(*R*) = 0.5, ⟨*z*⟩ = 0, var(*z*) ≈ 0.34. The remaining parameters are defined the same as the main text and taken to be: *α* = 1, *β* = 0.25, *λ* = 250, *λ*_leak_ = 0, *γ* = 1.5, *D* = 1, *κ* = 5, *M*_data_ = 500.

## Appendix C: Series solution for variance of population

In this section, we show the derivation of the variance result in Section III B. The calculations closely follow [63] on the stochastic cable equation. As noted in (12), the (2) SPDE can formally be solved by integrating the stochastic point source noise against the Green’s function *G*(*x, t*) that satisfies

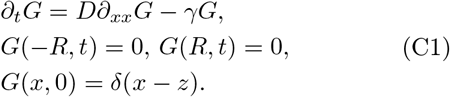

This Green’s function is [65]

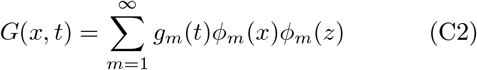

With

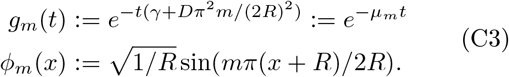

By (10), computation of the variance of *n* requires knowledge of the second moment of the underlying stochastic intensity. That is,

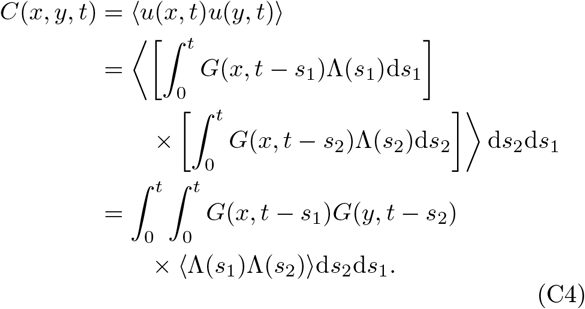

The autocorrelation of the dichotomous process is classical [60], and satisfies

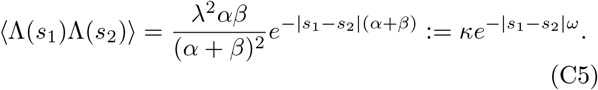

Ultimately, this gives us

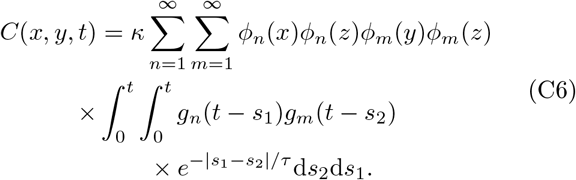

We defer evaluating this, noting that ultimately we want to spatially integrate *C*(*x, y, t*), and that

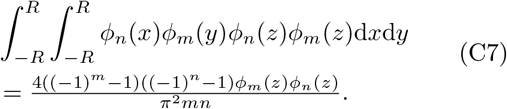

With some help from `Mathematica`, we can also integrate

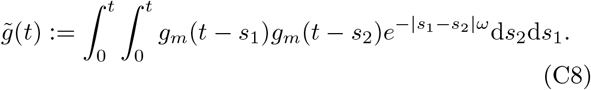

The expression is unwieldy, but after taking lim_*t* → ∞_, we get

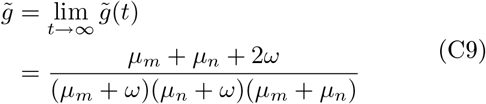

The temporal terms (C9) and spatial terms (C7) can be combined into (C6) to nearly give the infinite sum in the main text. The last ingredient is the observation that only odd terms of (C7) are nonzero, so we-reindex *m*→2*m* 1, *n*→2*n −* 1 and (11) is recovered. Figures with predictions from the series truncate after 1000 terms.

